# Hippocampal Activity Dynamics During Contextual Reward Association in Virtual Reality Place Conditioning

**DOI:** 10.1101/545608

**Authors:** Sidney B. Williams, Moises Arriaga, William W. Post, Akshata A. Korgaonkar, Jose A. Morón, Edward B. Han

**Affiliations:** Department of Anesthesiology, Washington University School of Medicine, St. Louis, MO, USA; Pain Center, Washington University School of Medicine, St. Louis, MO, USA; Department of Neuroscience, Washington University School of Medicine, St. Louis, MO, USA; Department of Psychiatry, Washington University School of Medicine, St. Louis, MO, USA

## Abstract

Exposure to environmental contexts associated with drug use can induce cravings that promote continued use and/or relapse. Opioid abuse is marked by high relapse rates, suggesting that contextual memories formed during opioid use may be particularly strong. While it is known that reward-seeking behavior is controlled by the mesolimbic reward circuit, little is understood about how contextual memories are altered by drug use. The dorsal hippocampus (dHPC) is necessary for multiple types of contextual learning and the place-specific activity of CA1 place cells map out space in a given environment. Here we examined the neuronal representation of context as animals developed morphine-paired environmental associations using a conditioned place preference (CPP) paradigm. To investigate changes in the hippocampal encoding before, during, and after drug-pairing, we developed a virtual reality (VR) morphine CPP (Mor-CPP) paradigm and used *in vivo* two-photon calcium imaging to record the activity of CA1 pyramidal neurons. We found increased activity in rewarded contexts following real-time operant conditioning with water rewards, but not after Mor-CPP training, suggesting different neural encoding mechanisms for natural reinforcers and morphine.

## INTRODUCTION

Associations between drug use and the drug administration environment, and/or drug-associated discrete cues, can persist throughout the stages of addiction, past initial drug-taking, to promote continued use and increase relapse (Daglish *et al.*, 2001; See, 2002). Reward seeking behavior is controlled by the mesolimbic reward circuit, in which the nucleus accumbens (NAc) integrates dopaminergic reinforcement signals from the VTA with environmental-specific stimuli encoded by glutamatergic inputs, largely from the ventral hippocampus (vHPC) (Phillips *et al*., 2003; Stuber *et al*., 2008 Britt *et al.* 2012). However, the vHPC gives a relatively coarse representation of space and is strongly associated with stress and emotional state, while, in contrast, the dorsal hippocampus (dHPC) is necessary for many types of contextual and spatial learning (Jung *et al*., 1994; Poucet *et al*., 1994; Moser *et al*., 1995; Moser and Moser, 1998; Fanselow and Dong, 2010). However, little is understood about its role in drug-context associations. Recent evidence suggests that the dHPC may play a critical role in forming these associations. AMPA and NMDA receptor-mediated plasticity in the dHPC are necessary for opioid-induced contextual learning, conditioned place preference (CPP) and context-dependent behavioral sensitization (Drake *et al.*, 2007; Zarrindast *et al*., 2007; Billa *et al.* 2009; Billa *et al*., 2010; Fakira *et al*., 2014; Portugal *et al*., 2014; Xia *et al*., 2011). Additionally, cocaine-CPP can be abolished by silencing dHPC CA1 pyramidal neurons active in the cocaine-paired environment (Trouche *et al*., 2016). While these studies suggest that the CA1 region of dHPC is important in the formation of reward-associated contextual representations for certain drugs of abuse, it is unknown specifically how opioid use might alter contextual representations in the hippocampus.

To address this question, we developed a virtual reality conditioned place preference (VR-CPP) behavioral paradigm using *in vivo* two-photon calcium imaging to record dHPC CA1 neuronal activity during the development of morphine-paired contextual associations. Two-photon imaging provides high-resolution activity recording from large cell populations in awake, behaving animals. We found that changes in hippocampal activity differed between Pavlovian conditioned morphine CPP (Mor-CPP) and operant contextual conditioning with a natural reinforcer (water). We hypothesize that the salience of opioids drives differential place representation in the dorsal hippocampus compared to natural rewards.

## METHODS

### Animals

All experiments were approved by the Washington University Animal Care and Use Committee. C57BL/6J mice ages 2-4 months of age were used for all experiments (Jackson Labs). Both male and female mice were used.

### Viral Injections and Hippocampal Window Implantation

Surgical and two-photon imaging procedures used were described previously, with minor modifications (Arriaga and Han, 2017). Mice were injected with AAV1.Syn.NES.jRGECO1a.WPRE.SV40 (Penn Vector Core, University of Pennsylvania) at 8-10 weeks of age. Mice were anesthetized with 1-3% isoflurane. The virus at a titer of 2.95 × 10^13^ g.c. was diluted 1:1 with PBS and pressure injected through a 0.5 mm diameter craniotomy above the left cortex through a micropipette targeted to the CA1 layer of the hippocampus (AP: −1.7 mm, ML: −1.6 mm, DV: −1.35 mm from bregma). The scalp incision was sealed with Metabond (Parkell) and a custom-cut titanium headplate (eMachineShop) was attached to the skull. Mice were water restricted (0.7-1 mL of water/day) to reduce their weight down to approximately 75% of their starting weight.

After 1 week, the headplate was removed, a larger craniotomy (2.8 mm) was made and the cortex above the dorsal hippocampus was aspirated. The imaging cannula [2.8 mm outer diameter, 2.36 mm inner diameter, 1.5 mm height (Microgroup); 2.5 mm round coverslip (Potomac Photonics)] was inserted at a 7-10° angle to match the angle of the dorsal side of the hippocampus and secured in place in the craniotomy above the hippocampus by Kwik-Sil elastomer (World Precision Instruments). Metabond darkened with carbon powder (Sigma-Aldrich) to prevent VR light from entering the microscope objective was applied to the skull around the cannula to close the incision and attach the headplate and light cup. Animals were allowed to recover for 2 weeks with continued water scheduling before VR training.

### VR Conditioned Place Preference

We designed a three chamber VR-CPP apparatus, similar to a classical CPP apparatus, composed of two conditioning chambers with distinct visual cues (vertical black and gray stripes vs. horizontally elongated polygons) connected by a third neutral white chamber. Mice were head fixed and allowed to freely run on a Styrofoam ball (Floracraft, 8 inch diameter) suspended by air pressure (Fig. 1). Movement of the ball was tracked by a computer mouse (Logitech), converted to forward and yaw velocities by custom written software in LabView, then fed into a custom virtual reality engine written in Matlab (ViRMEn; Aronov and Tank, 2014), which used the transformed movement information to continually update the visual display, permitting the animal to control navigation through the VR environment. The forward ball movement gain was set so 2.8 rotations of the ball, equivalent to 180 cm of distance traveled, traversed the long axis of the track. Yaw gain was set so 12 ball rotations equaled a 360° rotation in VR. The visual VR environment was back projected using two projectors (Optoma 750ST) onto a custom-made curved screen positioned 12 inches from the mouse encompassing 180° of horizontal (azimuth), and −16° below to +35° above horizon of the mouse. Water rewards (3-4 μL/reward) were controlled by a transistor-transistor logic (TTL) output from the VR engine.

**Figure 1.**
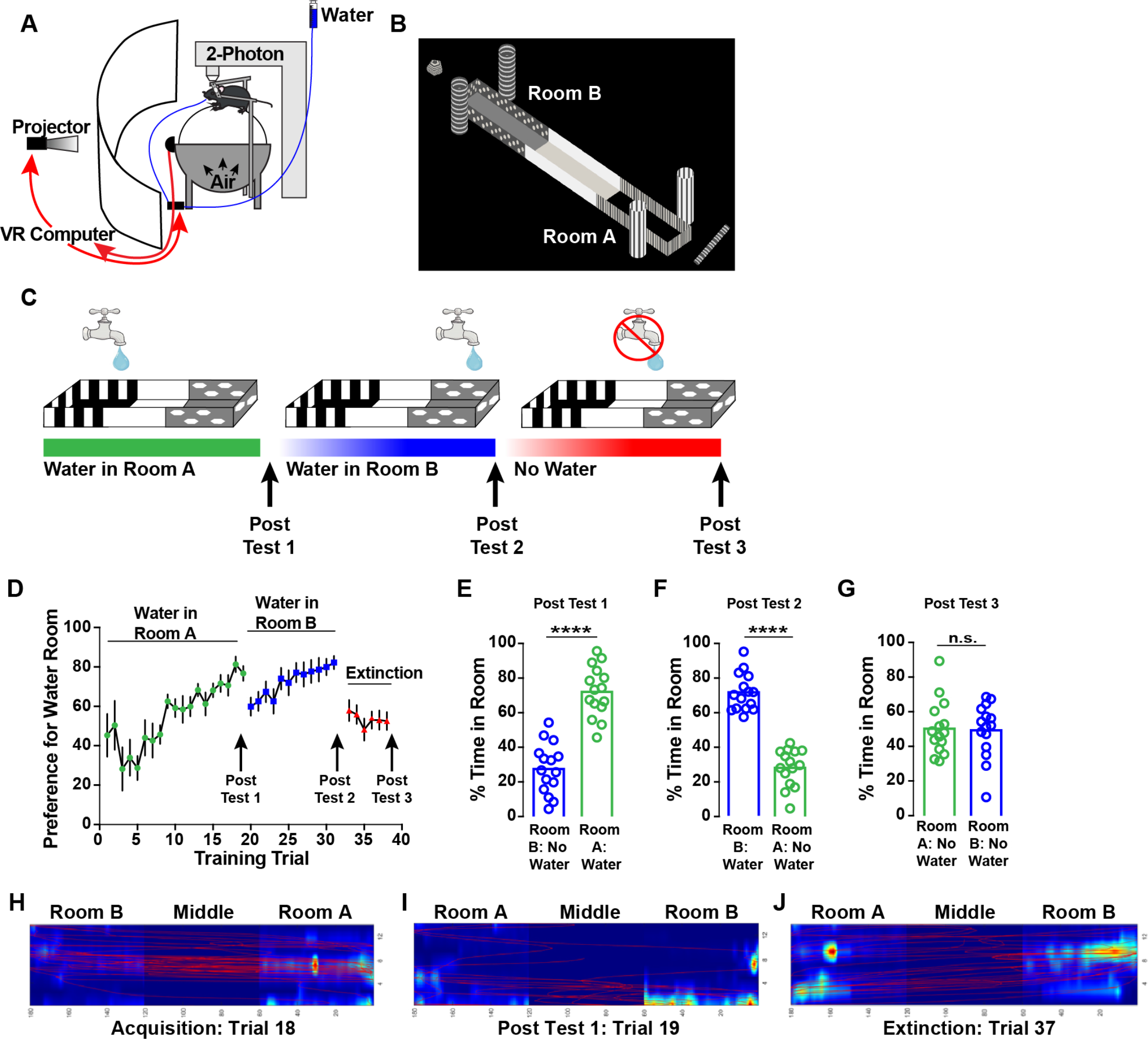
Virtual Reality (VR) Contextual Operant Conditioning with Water Rewards. A) Schematic of experimental approach. Mice are head-fixed and run on a spherical treadmill (Styrofoam ball) floating on a cushion of air. Ball movement is tracked by computer mouse and converted to forward and rotation velocities by custom written software (LabView) that feeds into the VR engine (ViRMEn, Matlab), which updates the projected visual scene. Water rewards are given at fixed time intervals for spending time in the rewarded room. B) View of VR-Conditioned Place Preference (CPP) environment. Mouse VR locomotion is restricted within the walls of the central corridor. C) Time course of water training and extinction. Animals were given two training trials per day and receive a water reward for spending 5-15 s continuously in the water-paired compartment. After criterion, room preference is tested with no rewards (Post Test 1). Then water rewards are switched to the opposite compartment, followed Post Test 2, forced abstinence extinction, with no rewards, and then Post Test 3. D) Real time place preference during operant conditioning for water rewards in VR. E) Post Test 1 after conditioning in Room A and F) Post Test 2, after switching water rewards to Room B.G) Forced abstinence in VR extinguishes preference for the water-paired compartment. H-J) Representative occupancy heat maps depicting time spent in the water-paired compartments and navigation path (red line) during trial 18 (last acquisition) (H), Post Test 1 (I), and trial 37 (last extinction trial) (J). Data are represented as mean +/− S.E.M. Significance was tested using a paired, two-tailed *t*-test: \*\*\*\**p* < 0.001, n.s. = not significant (*p* > 0.05); *n* = 15.

### Two-Photon Calcium Imaging

For imaging, we used a Neurolabware laser-scanning two-photon microscope with an electric tunable lens (ETL; Optotune, EL-10-30-NIR-LD) in the excitation light path, in combination with a f = −100mm offset lens. We sequentially imaged 4-6 axial planes spanning <180 μm in the *z*-axis at a total frame rate of 31 Hz with bidirectional line scanning. Emitted fluorescence was detected by a GaAsP photo-multiplier tube (Hamamatsu, 10770-40) after bandpass filtering (Semrock, BrightLine 607/70). Movies were collected through a 16×, 0.8 numerical aperture objective (Nikon, CFI 75 LWD 16X) with fields of view of ∼500 μm × 500 μm. Ti:sapphire laser (MaiTai, Spectra-Physics) light was tuned to 1015-1020 nm for jRGECO1a imaging. Laser power was ~25-50 mW after the objective and was set independently for each imaged plane using a pockels cell (ConOptics, 350-80-LA-02 KD, BK option, 302RM driver). Raw ball-speed data (forward and yaw, measured by computer mouse and exported via a National Instruments PCI-6221 data-acquisition board), VR data (track position, head direction, rewards, exported from the VR computer via a National Instruments PCI-6229 data-acquisition board), and imaging timing (each imaging frame was marked by a TTL output from the microscope-controller board) were synchronized by collecting all data in a single file using pClamp (RRID:SCR_011323) via a Digidata 1322 digitizer (Molecular Devices).

### Data Analysis and Statistics

Behavioral data, sampled at 1kHz, was downsampled to match the imaging frame rate. Residency in VR-CPP chambers was determined by the virtual reality position of the animals. The first 60cm in the long axis of the environment corresponded to chamber A and 121-180cm corresponded to chamber B. Conditioned place preference performance was assessed by comparing the ratio of time mice spent in the designated VR-CPP chamber to the total time the mouse spent in both conditioning chambers; time spent in the middle, neutral, chamber was not included.

Calcium imaging videos were processed, and cellular fluorescent time series were extracted using Suite2p (Pachitariu, 2017); the resulting fluorescent data were analyzed using custom written Matlab scripts. Slow fluctuations in fluorescence were removed from calculations of ΔF/F0 by calculating F0 using the eighth percentile of fluorescence intensity from a sliding window 30 s around each time point. Neuropil contamination was removed by subtracting a perisomatic fluorescence signal from an annulus between 5 and 20 μm from each ROI, excluding any other possible ROIs (F_Corrected-ROI_ =F_ROI_ – .8 * F_Neuropil_); (Peron et al., 2015, Dipoppa et al. 2018). ROIs were manually curated to eliminate putative interneurons based on morphology and firing patterns. Additionally, any cells with an activity skewness below 2 were classified as possible interneurons and excluded from analysis. The corrected fluorescence traces were subjected to analysis of the ratio of positive-to negative-deflecting transient size and durations. Significant transients were detected with 0.1% false positive error rates; these significant transients were used in the subsequent analysis. ΔF/F0 traces were set to 0 at all time points not identified as significant transient. Initial cellular preference for VR chamber was assessed using a cellular preference index measure, defined as the difference between the mean fluorescent activity in the morphine and saline associated chambers, normalized by the sum of these means (CPI = (μ_MOR_ – μ_SAL)/_ (μ_MOR_ + μ_SAL)_).

Statistical analysis was performed using Graph Pad Prism 6 software. For behavioral and imaging experiments, a paired two-tailed *t*-test was used to compare animal’s preferences pre- vs. post-conditioning. Wilcoxon rank test compared to 50% was also used to assess animal’s preference for the drug-paired compartment. Statistical significance was considered with a *p* < 0.05 in all experiments.

## RESULTS

### Establishment of a reward-associated VR-CPP paradigm

We designed a three chamber VR-CPP apparatus, based on the classical CPP apparatus (Tzschentke,1998), composed of two conditioning chambers with distinct visual cues (vertical black and gray stripes vs. horizontally elongated polygons) connected by a third neutral white chamber (Fig. 1B). Distinct distal cues were placed outside the walls to aid animal orienting. Mice were water scheduled to motivate them to seek water rewards. Animals underwent two operant conditioning training trials (21.5 min each) per day during which they received a water reward for spending 5-15 seconds continuously in the water-paired compartment of the VR environment, whereas no reward was given in the opposite chamber (Fig. 1C). Within 8-12 days the animals reached the learning criterion (>70% of time spent in the water-paired compartment for 4 consecutive trials) (Fig. 1D). After reaching criterion, mice were given a post-conditioning test (Post Test 1) in which they had full access to the VR environment without access to water rewards. During Post-Test 1, mice demonstrated a significant preference for the water-paired chamber even though there was no water reward available (*n* = 15, 27.72 ± 3.76 % time in non-water-paired room, 72.28 ± 3.76 % time in water-paired room; two-tailed paired *t*-test: *p* < 0.0001) indicating contextual memory-based reward seeking behavior. To ensure that no procedural bias was created, reward contingency was then switched to the opposite compartment (Fig. 1D). Mice reached the learning criterion faster in the second water-paired room compared to the first, consistent with typical reversal learning (Winocur and Olds, 1978). When tested during a second post-conditioning test (Post Test 2), mice demonstrated a significant preference for the new water-paired chamber (*n* = 15, 28.41 ± 2.78 % time in non-water-paired room, 71.59 ± 2.78 % time in water-paired room; two-tailed paired *t*-test: *p* < 0.0001). After undergoing an extinction protocol, in which animals were exposed to the VR-CPP environment without rewards, mice demonstrated extinguished reward-context associations (Fig. 1C). The extinction post test (Post Test revealed that the prior behavior preference was eliminated (*n* = 15, 50.46 ± 4.05 % time in Room A, 49.54 ± 4.05 % in Room B; two-tailed paired *t*-test: *p* = 0.91). Altogether, our data validate the development of an operant context-dependent paradigm to study natural reward seeking behavior in mice. The use of VR technology enables the construction of multiple behavioral environments with precise control over all salient sensory stimuli to dissect the contribution of individual cues and contextual variables. Finally, in contrast to prior studies, here we are able to use the same apparatus to examine contingent and non-contingent behavior.

Following operant water-conditioning, mice underwent a Mor-CPP protocol in the VR environment (Fig. 2A), modified based on our previous reports (Portugal *et al*., 2014; Fakira *et al*., 2016). First, animals underwent a pre-conditioning test in which they ran freely, without reward, to establish a baseline for time spent in each compartment as a measure of any inherent bias individual animals may have for one context over the other. During conditioning, morphine was paired with the non-preferred chamber for each mouse (Tzschentke, 1998). The following day, animals began morphine contextual conditioning, as depicted in Fig. 2. During morning training sessions, animals received a saline injection (Sal; 15 mg/kg, s.c.) and restricted to one chamber in the VR-CPP environment by an additional virtual wall. In the afternoon, animals received a morphine injection (15 mg/kg, s.c.) and restricted to the opposite chamber. Animals received 8 days of saline/morphine conditioning before they underwent a post-conditioning preference test. No vehicle or drug injection was administered prior to the post-conditioning test and animals were given free access to the entire VR environment.

**Figure 2.**
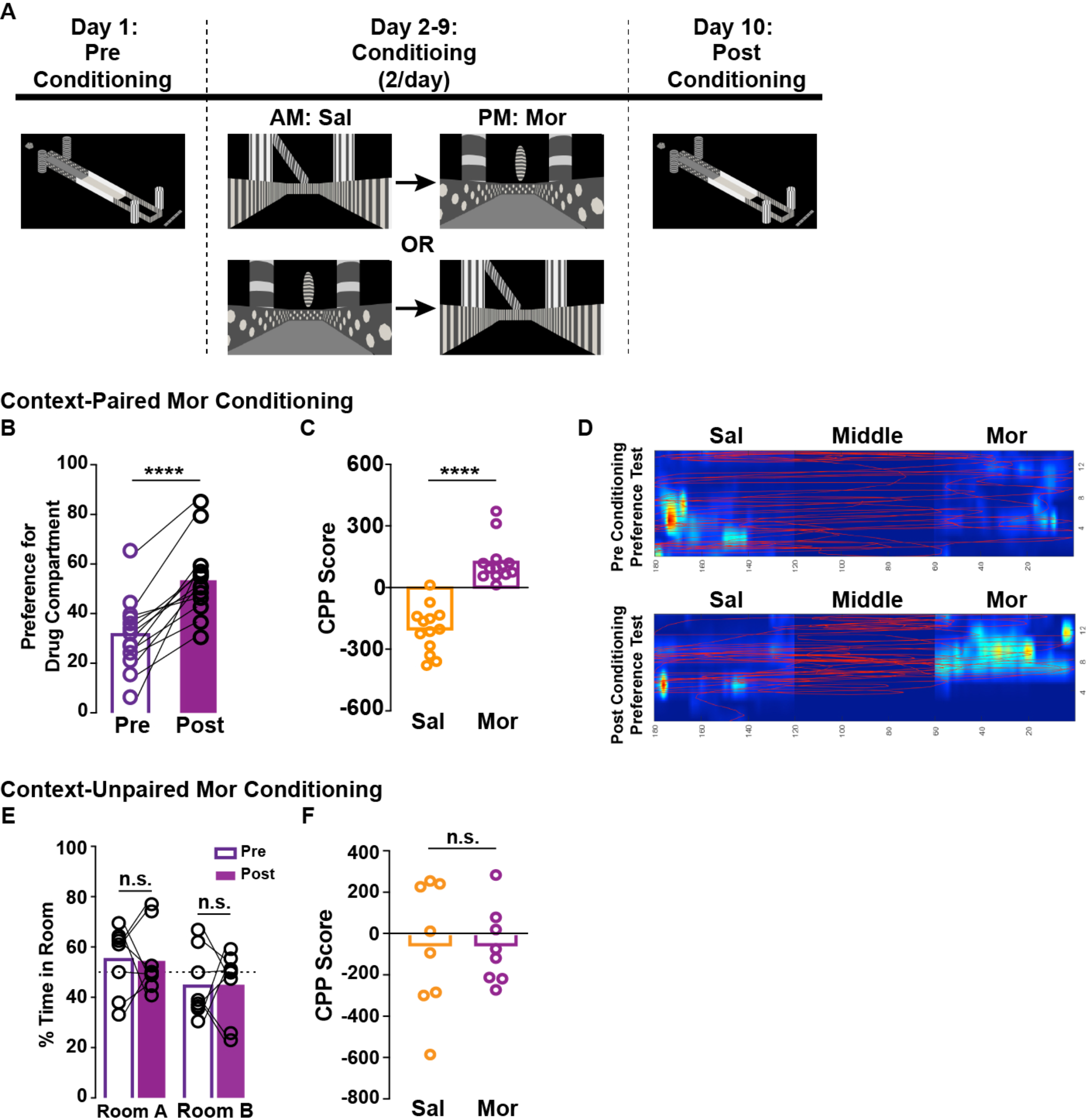
Morphine Conditioned Place Preference (Mor-CPP) in VR. A) Schematic representation of the behavioral paradigm used for Mor (15mg/kg) conditioning in VR. For conditioning sessions, animals are restricted to one room by addition of a virtual wall. For context-paired Mor conditioning, the animals are administered Mor in the same chamber every day. In contextually unpaired Mor conditioning, the animal is administered Mor in alternating rooms. B) Behavioral preference for the Mor-paired compartment demonstrated in percent time (in the absence of drug). C) CPP score (post – pre conditioning time) for the Sal-paired and Mor-paired compartments. Mice demonstrate a switch in place preference for the Mor-paired compartment following contextual Mor conditioning. *n* = 13; Paired, two-tailed *t*-test; \*\*\*\**p* < 0.001. D) Representative occupancy heat maps depicting time spent in the Mor-paired and Sal-paired compartments and navigation path (red line) during the pre-conditioning preference test (top) and the post-conditioning preference test (bottom). E) Histogram of time spent in each room during pre- and post-conditioning. Contextually unpaired Mor conditioning does not illicit a behavioral preference. F) CPP score for unpaired Mor-conditioning. Data are represented as mean +/− S.E.M. Significance was tested using a paired, two-tailed *t*-test: *p* > 0.05; *n* = 9.

During the post-conditioning test following morphine-paired contextual conditioning, we observed a significant switch in preference for the Mor-paired VR context (Fig. 2B; n = 13, 31.67 ± 4.04 % time in Sal-paired room, 53.66 ± 4.2 % time in Mor-paired room; two-tailed paired *t*-test: *p* < 0.0001). To rule out any effects of the morphine treatment itself, we conducted additional unpaired morphine-conditioning in which morphine is given daily, but in alternating compartments (as we have previously described in Portugal *et al*., 2014; Fakira *et al*., 2016). On average, animals that underwent unpaired morphine-contextual conditioning showed no preference for either compartment, suggesting that the preference observed in the morphine-paired VR-CPP paradigm is independent of morphine administration *per se*, but rather depends on the formation of drug-induced contextual memories (Fig. 2; *n* = 8, 2way ANOVA: *F*(1,14) = 0.02, *p* = 0.88).

Here we provide evidence that VR environments are sufficient to establish opioid-induced contextual associations. The behavioral preference observed in the VR-CPP is similar to that generated using classical CPP approaches. While we have successfully demonstrated that virtual environments are sufficient to induce conditioned place preference behavior, the temporal dynamics of the neuronal plasticity underlying the formation and maintenance of these contextual drug-paired associations remains elusive. The VR-CPP allows, for the first time, the combination of drug-paired contextual behavioral paradigms with *in vivo*, high resolution two-photon imaging to examine the cellular activity mediating this behavior in real-time.

### Neurons are more active in water-paired contexts after operant contextual conditioning, but not in non-contingent Mor-paired environments

To analyze neuronal activity in dHPC CA1 during reward-paired contextual associations, we combined two-photon Ca^2+^ imaging with our VR-CPP behavioral paradigm. Cells were infected with AAV1.Syn.NES.jRGECO1a.WPRE.SV40 and calcium activity was imaged in multiple z-planes of CA1 at a frame rate of 5.2 – 7.7 Hz per plane (Arriaga and Han, 2017) (Fig. 3A, *N* = 8 mice, 444 ± 229 neurons per mouse, per imaging session). Mice were imaged on set days throughout the operant water conditioning, extinction, and subsequent Mor-CPP. We found that over the course of initial operant water conditioning, pyramidal cells showed preferential activity in the water-paired chamber, as calculated by cell preference ratios [neuronal activity correlated to time spent in one contextual compartment vs the other; (μ_A_-μ_B_)/ (μ_A_+ μ_B_)], compared to the non-paired chamber by Day 10 (Fig. 3C and D; *N* = 8, two-tailed paired *t*-test: Day 1 *p* = 0.4; Day 5 *p* = 0.33, Day 10 *p* = 0.005), although this preference did not persist through the post-conditioning test (Post Test 1), where no water rewards were available (Fig. 3E; *N* = 8, two-tailed paired *t*-test: Post-conditioning Test *p* = 0.06). When water rewards were switched to the other compartment (Fig. 3F), we also observed the development of robust preferential activity for the new rewarded context (Fig. 3G; *N* = 5-9, two-tailed paired *t*-test: Day 1 *p* = 0.55; Day 5 *p* < 0.0001, Day 10 *p* = 0.001). This significant population preference persisted throughout the post-conditioning test (Post Test 2; Fig. 3H; *N* = 8, two-tailed paired *t*-test: Post-conditioning Test *p* = 0.0002). The population activity preference accompanying operant water-reward place preference was eliminated during extinction training (Fig. 3I-K; *N* = 9, two-tailed paired *t*-test: *p* = 0.0011).

**Figure 3.**
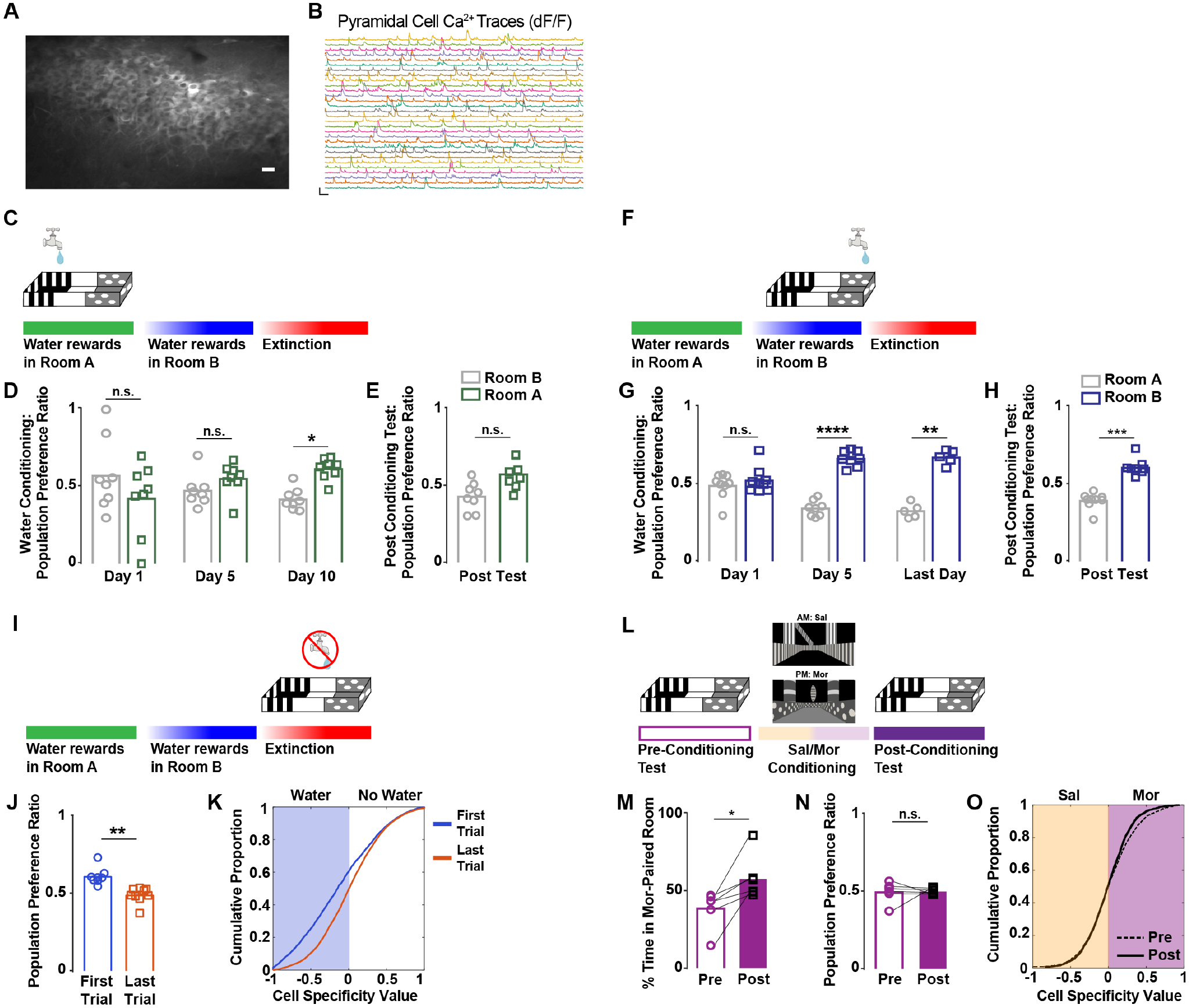
CA1 pyramidal neurons more active in water-paired context after real-time operant conditioning, but not after Mor-CPP. A) Left: AAV1.Syn.NES.jRGECO1a.WPRE.SV40 infected CA1 pyramidal neurons imaged in awake, behaving mice (Scale bar = 20 μm). B) Calcium transient traces from 30 pyramidal cells during VR-CPP conditioning in behaving animal (Scale bars: y = 1 dF/F, × = 10 sec). C) Schematic of water training in Room A. D) Pyramidal cell population preference ratios for Day 1, Day 5, and Day 10 of operant water conditioning. Population preference ratio is calculated by (μ_A_-μ_B_)/ (μ_A_+ μ_B_). E) Pyramidal cell population preference ratios for Room A during Post Test 1. F) Schematic of water training in Room B. G) Pyramidal cell population preference ratios for Day 1, Day 5, and Last Day of operant water conditioning in Room B. H) Pyramidal cell population preference ratios for Room B Post Test (2). I) Schematic of extinction of water training, following water-paired rewards in Room B. J) Pyramidal cell population preference ratios the first and last day of extinction training. K) Cumulative proportion of cell specificity values, as calculated by (μ_A_-μ_B_)/ (μ_A_+ μ_B_). The difference between mean ΔF/F in each region, normalized by the sum of mean ΔF/F. L) Schematic representation of the behavioral paradigm used for Mor (15mg/kg) conditioning in VR. M) Behavioral preference for Mor-paired compartment represented by percent time. N) Pyramidal cell population preference ratios before and after Mor conditioning. O) Cumulative proportion of cell specificity values, as calculated by (μ_A_-μ_B_)/ (μ_A_+ μ_B_). The difference between mean ΔF/F in each region, normalized by the sum of mean dF/F. Data are represented as mean +/− S.E.M. Significance was tested using a paired, two-tailed *t*-test with Bonferroni correction for multiple comparisons; \**p* < 0.05, \*\**p* < 0.01, \*\*\**p* < 0.001, \*\*\*\**p* < 0.0001, n.s. > 0.05; n = 5-9.

To determine whether this population activity preference was mediated by the VR operant learning paradigm or the reward itself, we performed the same population imaging analysis during our Mor VR-CPP behavioral paradigm, as described in Fig 2. Although animals show a significant switch in behavioral preference for the Mor-paired room during the post-conditioning test (Fig. 3M; *N* = 6, two-tailed paired *t*-test: Pre- vs. Post-conditioning Test *p* = 0. 02), we did not observe a corresponding pyramidal cell population activity preference (Fig. 3N-O; *N* = 6, two-tailed paired *t*-test: Pre- vs. Post-conditioning Test *p* = 0. 88). These results suggest that the hippocampal representation of a natural reward-associated place differs from a non-contingent drug-reward associated context. It is also possible that these differences arise because the water VR-CPP was conducted as an operant paradigm in contrast to the Mor VR-CPP, which was conducted following Pavlovian conditioning in which morphine was administered in a non-contingent manner. Future analysis will focus on how hippocampal representations of place differ between reward-context associations driven by a natural reinforcer and drugs of abuse. Finally, we propose that the long-lasting association between opioid use and environmental context arises, in part, from a fundamental alteration in place coding by pyramidal neurons in the dorsal hippocampus.

Here we present a powerful and flexible virtual environment for associating rewards with context. This platform, in combination with two-photon imaging, allows us to monitor the activity of large neuronal ensembles during the formation of reward-associated memories. This approach will allow us to longitudinally follow the same cells over time (Ziv *et al.,* 2013; Driscoll *et al.,* 2017; Xia *et al.,* 2017), correlate activity across multiple brain areas (Stirman *et al.,* 2016; Sofroniew *et al.,* 2016; Sjulson *et al.,* 2018), track structural plasticity (Trachtenberg *et al.,* 2002; Grutzendler *et al.,* 2002; Fakira *et al.,* 2016), and record from genetically identified neuronal populations (Arriaga and Han, 2017) to provide a comprehensive picture of how pathological memories of drug-associated contexts form and persist. This information may give rise to new targeted strategies for breaking the cycle of relapse that perpetuates the current opioid epidemic.

## Acknowledgements

This work was supported with funding from the National Institutes of Health (NIH) through a Cutting-Edge Basic Research Award (CEBRA-DA042581) and the Washington University, Department of Psychiatry, Addiction Training Grant (T32-DA007261).

